# Inverted Regulation of Multidrug Efflux Pumps, Acid Resistance and Porins in Benzoate-Evolved *Escherichia coli* K-12

**DOI:** 10.1101/531178

**Authors:** Jeremy P. Moore, Haofan Li, Morgan L. Engmann, Katarina M. Bischof, Karina S. Kunka, Mary E. Harris, Anna C. Tancredi, Frederick S. Ditmars, Preston J. Basting, Nadja S. George, Arvind A. Bhagwat, Joan L. Slonczewski

## Abstract

Benzoic acid, a partial uncoupler of the proton motive force (PMF), selects for sensitivity to chloramphenicol and tetracycline during experimental evolution of *Escherichia coli* K-12. Transcriptomes of *E. coli* isolates evolved with benzoate showed reversal of benzoate-dependent regulation, including downregulation of multi-drug efflux pumps; the Gad acid resistance regulon; the nitrate reductase *narHJ*; and the acid-consuming hydrogenase Hyd-3. However, the benzoate-evolved strains had <u>increased</u> expression of OmpF and other large-hole porins that admit fermentable substrates and antibiotics. Candidate genes identified from benzoate-evolved strains were tested for their roles in benzoate tolerance and in chloramphenicol sensitivity. Benzoate or salicylate tolerance was increased by deletion of the Gad activator *ariR*, or of the *slp-gadX* acid fitness island encoding Gad regulators and the multi-drug pump *mdtEF*. Benzoate tolerance was also increased by deletion of multi-drug component *emrA*, RpoS post-transcriptional regulator *cspC*, adenosine deaminase *add*, hydrogenase *hyc* (Hyd-3), and the RNA chaperone/DNA-binding regulator *hfq*. Chloramphenicol resistance was decreased by mutations in global regulators, such as RNA polymerase alpha-subunit *rpoA*, Mar activator *rob*,and *hfq*. Deletion of lipopolysaccharide biosynthetic kinase *rfaY* decreased the rate of growth in chloramphenicol. Isolates from experimental evolution with benzoate had many mutations affecting aromatic biosynthesis and catabolism such as *aroF* (tyrosine biosynthesis) and *apt* (adenine phosphoribosyltransferase). Overall, benzoate or salicylate exposure selects for loss of multi-drug efflux pumps and of hydrogenases that generate a futile cycle of PMF; and upregulates porins that admit fermentable nutrients and antibiotics.

**IMPORTANCE:** Benzoic acid is a common food preservative, and salicylic acid (2-hydroxybenzoic acid) is the active form of aspirin. At high concentration, benzoic acid conducts a proton across the membrane, depleting the proton motive force. In the absence of antibiotics, benzoate exposure selects against proton-driven multidrug efflux pumps and upregulates porins that admit fermentable substrates but also allow entry of antibiotics. Thus, evolution with benzoate, and related molecules such as salicylates, requires a tradeoff for antibiotic sensitivity—a tradeoff that could help define a stable gut microbiome. Benzoate and salicylate are naturally occurring plant signal molecules that may modulate the microbiomes of plants and animal digestive tracts so as to favor fermenters and exclude drug-resistant pathogens.

## INTRODUCTION

*Escherichia coli* and other enteric bacteria face high concentrations of organic acids, such as short-chain fatty acids that permeate bacterial cell membranes and acidify the cytoplasm (1, 2) and drive the accumulation of toxic anions (3, 4). Acids that cross the membrane in the unprotonated form can uncouple the proton motive force (PMF) (5–8). Benzoic acid and salicylic acid (2-hydroxybenzoic acid, the active form of aspirin) act as permeant acids and as partial uncouplers. These molecules are abundant in in the plant rhizosphere (9) and in human diets in the form of food preservatives, pharmaceutical products, and natural plant secondary metabolites (10–13). We are investigating the molecular effects of benzoate derivatives on bacteria.

Benzoate and salicylate induce low-level resistance to antibiotics, via the Mar regulon (14) as well as mar-independent pathways that are poorly understood (15). The Mar regulon intersects with the Gad acid resistance regulon (16, 17) which includes a major region of acid-stress regulators and multidrug efflux pumps (MDR) (**Fig. 1**). Surprisingly, however, experimental evolution in the presence of benzoate leads to deletion of acid resistance systems, and decreased resistance to the antibiotics tetracycline and chloramphenicol (18). Isolates from populations serially cultured for 2,000 generations show loss of MDR pumps and regulators such as *emrA, emrY*, and *marRAB*. Similarly, experimental evolution in the presence of the strong uncoupler, carbonyl cyanide m-chlorophenyl hydrazone (CCCP), yields isolates that have lost MDR pumps and regulators, with the exception of the EmrAB-TolC pump which directly exports CCCP (19, 20). These results suggest a hypothesis that aromatic acid uncouplers can amplify the fitness cost of MDR in the absence of antibiotics. The concept of reversing antibiotic resistance is of great interest for the gut microbiome as well as contaminated environments, where antibiotic-resistant strains may show a minimum selective concentration (MSC) as much as 100-fold lower than the minimum inhibitory concentration (MIC) (21).

**Figure 1.**
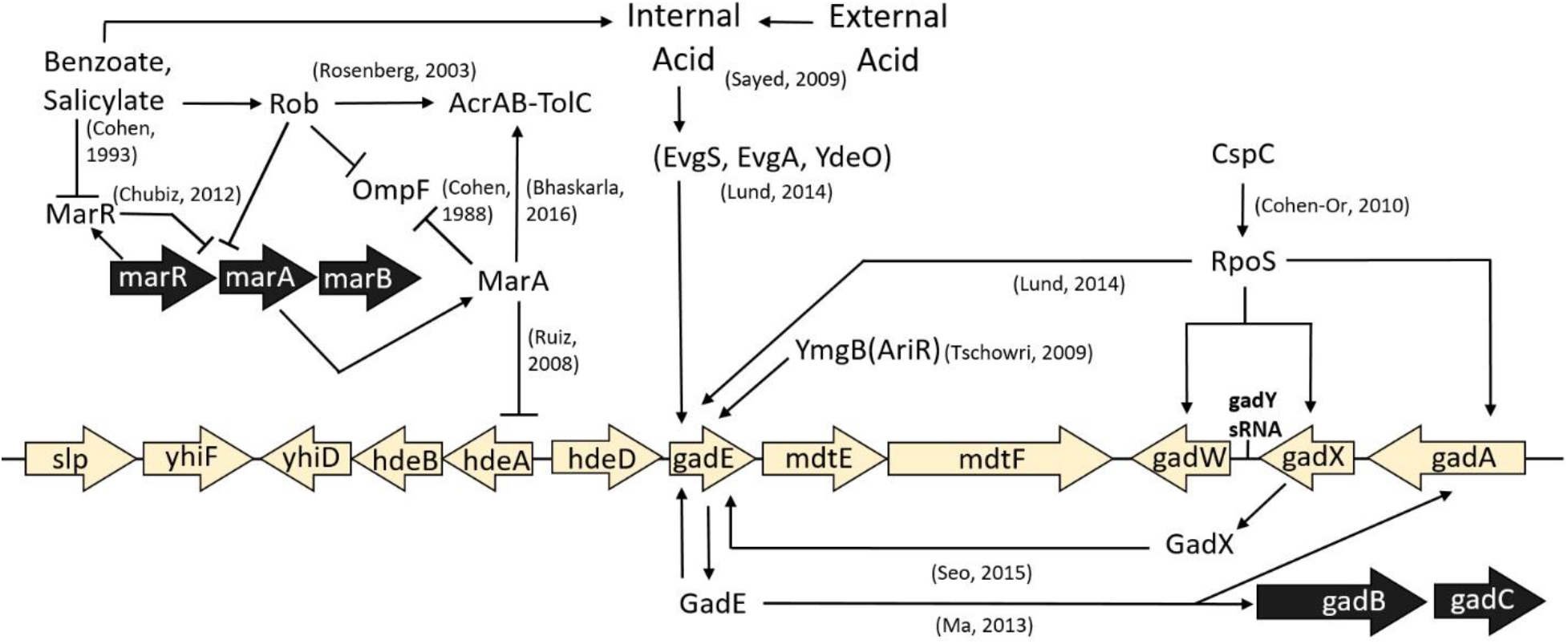
The Gad acid resistance regulon intersects with the Mar drug-resistance regulon. Selected components relevant to this work are shown.

We considered several mechanisms that might explain benzoate selection against drug resistance. When a partial uncoupler depletes PMF, the cell incurs energy stress, which selects against MDR pumps that spend PMF. Additionally, constitutive induction of stress response pathways carries a heavy energetic penalty; thus, deletion or downregulation of constitutively expressed pathways confers an energetic advantage. The penalty could arise from the cost of protein production, which increases non-linearly with additional transcription (22). In the case of acid stress, experimental evolution leads to loss of three acid-inducible amino-acid decarboxylase systems (23, 24). Also, PMF expenditure in itself has a fitness cost, as in the *lac* operon, where the major fitness cost of expression is the activity of LacY permease, driven by PMF (25). The depletion of PMF might favor expression of large-hole porins that increase uptake of fermentable substrates, with the tradeoff of increased uptake of antibiotics (26–28).

Most benzoate-tolerant isolates from the evolution experiment (18) acquired a mutation affecting the *slp-gad* (Gad) acid fitness island (16, 29–31). The Gad island includes acid-resistance regulators *gadE, gadW, gadX*(32); periplasmic acid chaperones *hdeA* and *hdeB*; and the MDR efflux components *mdtE, mdtF* (33, 34) (**Fig. 1**). The major regulator GadE upregulates several acid-tolerance genes, including two glutamate decarboxylase isoforms (*gadA*and *gadB*), whose activity increases cytoplasmic pH by consuming protons via decarboxylation of glutamate (16, 31, 35, 36). The MdtEF-TolC complex exports antibiotics from the cytoplasm, driven by PMF. Our benzoate-evolved isolates (18) have acquired point mutations or deletions throughout *slp-gad*, as well as knock-out or point mutations in *emrA*, the membrane-fusion protein component of the EmrAB-TolC MDR pump (37, 38); *mdtA* (39, 40); and *emrY* (41). Other mutations affect MDR regulators such as *cpxA* (42), *ariR* (43), *arcA* (44), and *rob* (45).

In one population, exposure to benzoate has selected for deletion of *marRAB*, a multidrug resistance operon which is induced by MarR binding salicylate or benzoate (15, 45). MarRAB has homologs throughout bacteria and archaea, including many drug-resistant clinical isolates (46). In *E. coli*, salicylate or benzoate relieves repression of MarA, which regulates over 60 genes involved in antibiotic resistance such as AcrAB-TolC MDR pump (14, 47) as well as *hdeAB* within the *slp-gad* island (17). MarA downregulates the large porin OmpF, which admits nutrients and antibiotics (48). Many targets of MarA are also subject to MarA homolog Rob (45, 46), which had a mutation in one benzoate-evolved isolate (18). Additionally, one strain acquired a mutation in the alpha subunit of RNA-polymerase which could affect the regulation of a wide array of genes. RNAP mutations lead to unexpected phenotypes, such as the downregulation of arginine decarboxylase caused by an *rpoC* mutation in an acid-evolved strain (23).

We sought to reveal the genetic mechanisms of benzoate tolerance found in the 2,000-generation strains, and to determine how these mechanisms intersect with antibiotic resistance. Here we report transcriptomic analysis of four 2,000-generation strains that showed a surprising long-term reversal of the short-term benzoate stress response. To dissect this response, and the mechanisms of antibiotic resistance reversal, we resequenced earlier populations from the benzoate evolution experiment with fewer mutations so as to reveal the order in which mutations were acquired. We report the effects of several candidate gene deletions and mutations on fitness in the presence of benzoate and of the antibiotic chloramphenicol. Our findings indicate several processes that mediate the antibiotic sensitivity of benzoate-evolved strains.

## RESULTS

### Early benzoate-selected mutations affect Gad, Mar and aromatic metabolism

The large number of mutations in the 2,000 generation strains made it difficult to assess which genetic changes were most likely to affect benzoate tolerance (18). As such, we decided to isolate and sequence strains from earlier generations that were more likely to contain fewer mutations. We isolated clones from frozen populations ancestral to those published for generation 2,000 (**Table 1**). Colonies were obtained on LBK agar from frozen populations corresponding to generations 900 and 1,400, for the microplate well populations A1, A5, C3, and G5. These populations at generation 2,000 had produced isolates A1-1, A5-1, C3-1, G5-1, and G5-2; these key strains are benzoate-tolerant, and all except G5-1 are more sensitive to chloramphenicol than the ancestral strain W3110 (18). Selected strains were sequenced (**Table S1**), and mutations were detected using *breseq* (49). Mutations for selected strains are listed in **Table 2**, alongside the mutations from generation 2,000 clones (18). All mutations for all clones sequenced are compiled in **Table S2**.

**Table 1.** Strains of *Escherichia coli* used in this study^1^

| Name | Genotype | Source |
| --- | --- | --- |
| W3110 | *Escherichia coli* K-12 | (110) |
| JLSK0001 | W3110 benzoate-evolved A1-1 | (18) |
| JLSK0014 | W3110 benzoate-evolved C3-1 | (18) |
| JLSK0030 | W3110 benzoate-evolved G5-1 | (18) |
| JLSK0031 | W3110 benzoate-evolved G5-2 | (18) |
| JLS0903 | W3110 Δ*hyaB::kanR* | (78) |
| JLS0905 | W3110 Δ*hycE::kanR* | (78) |
| JLS1010 | W3110 Δ*mdtE::kanR* | This work |
| JLS1025 | W3110 Δ*mdtF::kanR* | This work |
| JLS1517 | W3110 Δ*gadX::kanR* | This work |
| JLS1631 | W3110 Δ*cspC::kanR* | This work |
| JLS1632 | W3110 Δ*gadE::kanR* | This work |
| JLS1633 | W3110 Δ*rfaY::kanR* | This work |
| JLS1706 | W3110 Δ*emrA::kanR* | (20) |
| JLS1732 | W3110 $\Delta$(*slp-gadX*) | This work |
| JLS1771 | W3110 $\Delta$(*slp-gadX*) Δ*ariR::kanR* | This work |
| JLS1777 | W3110 $\Delta cspC::kanR \Delta rfaY::frt$ | This work |
| JLS1789 | W3110 $\Delta add::kanR$ | This work |
| JLS1820 | W3110 $\Delta rob::kanR$ | This work |
| JLS1823 | W3110 $\Delta ariR::kanR$ | This work |
| JLS1828 | W3110 $\Delta hycF::kanR$ | This work |
| JLS1616 | A1-1 $\Delta yhdN::kanR rpoA+$ | This work |
| JLS1617 | A1-1 $\Delta yhdN::frt$ | This work |
| JLS1791 | W3110 $\Delta hfq::kanR$ | This work |
| JLS1908 | W3110 $\Delta yhdN::kanR$ | This work |
| JLS1909 | A1-1 $\Delta yhdN::kanR$ | This work |

<sup>1</sup>Newly isolated strains from benzoate-evolved populations (18) are listed in Supplementary

The early-generation clones (900 or 1,400) were tested for adaptation to growth in 20 mM benzoate (**Fig. 2**; for all eight individual replicate curves, see **Fig. S1**). At this benzoate concentration, the ancestral strain W3110 stops growing and enters death phase by 12 h, whereas the 2,000-generation evolved strains grow more rapidly and reach stationary phase at about OD_600_ = 0.25 (18). All the A1 population isolates from generation 900 or 1,400 grew in 20 mM benzoate to a stable stationary phase. From the A5 population, only the A5-5 strain (generation 1,400) achieved a sustained endpoint. In population G5, all the early isolates grew well (G5-3, G5-4, G5-5); but in population C3, the generation 900 strains grew only slightly better than W3110. Overall, the various early-generation isolates showed differing rates of adaptation to benzoate.

**Figure 2.**
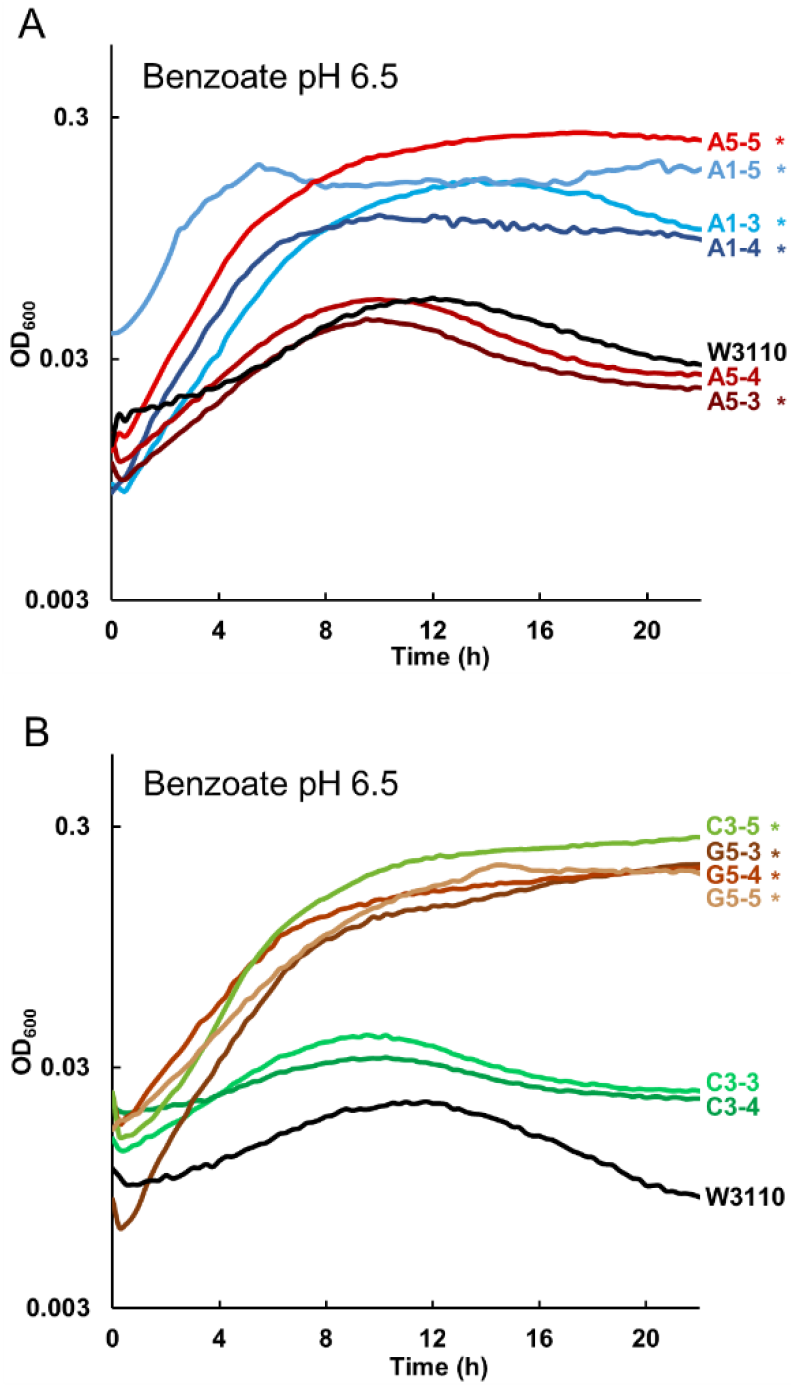
Isolates from early generations (500-1500 doublings) from populations A1, A5, C3, and G5. Strains were cultured in LBK 100 mM PIPES pH 6.5 with 20 mM sodium benzoate, as described under Methods. For each strain, the curve shown represents median value of OD_600_ at 16 h. Asterisk indicates significant difference from W3110, Tukey’s test p ≤ 0.05, n=4. All four replicates are shown in **Figure S2. A.** A5-5, A1-5, A1-3, and A1-4 outgrew ancestor W3110. A5-3 and A5-4 entered death phase after 12 h, as did ancestor W3110. **B.** C3-5, G5-3, G5-4, and G5-5 showed increased benzoate resistance compared to ancestor W3110. C3-3 and C3-4 entered death phase after 12 h, as did ancestor W3110.

Clones frozen before generation 1,000 had several mutations that persisted in the 2,000-generation strains (**Table 2**). Three of these mutations affected Gad-island regulation: the partial Gad island deletion *ΔmdtE-slp* (all isolates from population A1), the Gad activator *ariR* (population A1) (50), and GadE-activator *gadX* (population A5). In fact, more than half the isolates sequenced had one of seven different mutations within the *slp-gad* island (**Table S1**). Large deletions were likely mediated by upstream insertion sequences, a common finding in stress evolution experiments (18, 23). Thus our early-generation sequences confirmed that loss of Gad acid resistance was strongly selected by benzoate.

**Table 2.**
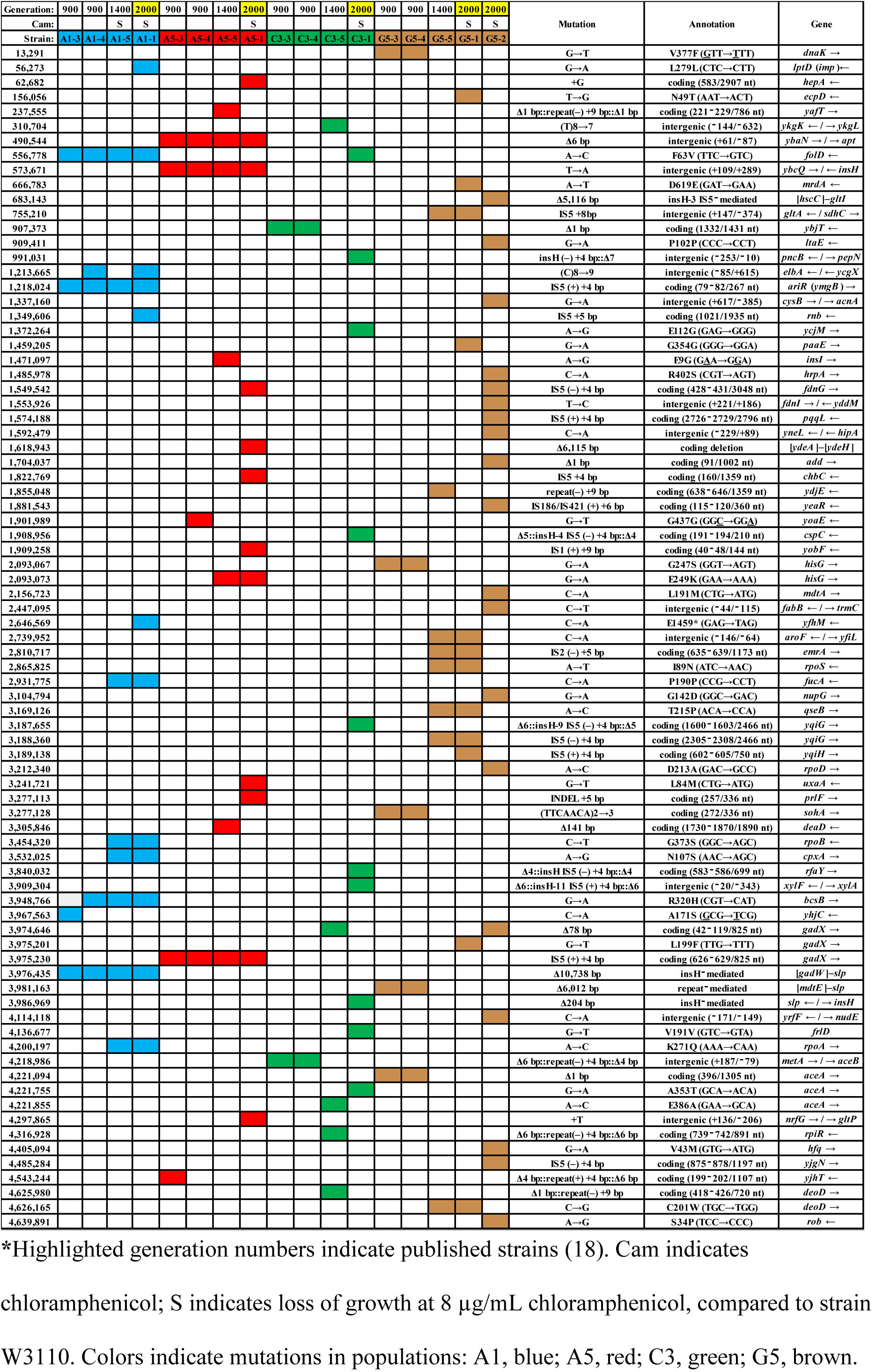
Mutations found in selected benzoate-evolved strains*

Loss of the Gad regulon, either by deletion or downregulation, includes loss of the MdtEF-TolC efflux pump, which is specifically upregulated by GadX (34). MdtEF-TolC exports a variety of antibiotics and toxic metabolites, including chloramphenicol (44, 51, 52). It is one of a number of MDR pumps and regulators reported to be lost or mutated in genomes evolved with benzoate (18) or with CCCP (20). Note that while *mdtEF* mutations did appear early, sensitivity to chloramphenicol was not detected before generation 1,400 or 2,000 (**Table 2**). This suggests that multiple mutations accumulate over generations contributing to the phenotype.

Various isolates had mutations affecting metabolism of nucleotides and aromatic amino acids; that is, structures with similarity to benzoate or salicylate (**Table S2**). Genes with early, persistent mutations include *folD* (5,10-methylene-tetrahydrofolate dehydrogenase/cyclohydrolase, for thymidine biosynthesis) (53) in population A1; and upstream of *apt* (adenosine phosphoribosyltransferase, for purine salvage) (54) in population A5. The 6-bp insertion upstream of *apt* was found by generation 500 in strain A5-6 and persisted through generation 2,000; and a frameshift in *apt* coding sequence was in strain E1-5 (**Table S2**). Strain E1-5, generation 1,000, had a mutation in *yeaS* (*leuE*) whose product effluxes leucine and toxic analogues. Biosynthetic genes with mutations included *aroF* (3-deoxy-d-arabino-heptulosonate 7-phosphate synthase; first step of tyrosine biosynthesis) (55) and *hisG* (ATP-phosphoribosyltransferase, first step of histidine biosynthesis) (56). Other mutations affected genes for aromatic catabolism and salvage: *add* (adenosine deaminase) (57); *deoD* (PNP, purine nucleoside phosphorylase) (58); *rihA* (ribonucleoside hydrolase) (59); *paaE* (phenylacetate degradation) (60); and *nupG* (nucleoside uptake transporter, PMF-driven) (61). These mutations in aromatic metabolism may represent responses to the benzoate uptake in the cytoplasm. Their effects might be associated with the induction of Mar regulon by intracellular aromatic intermediates that retard growth (62, 63).

### Transcriptomes of benzoate-evolved isolates show reversal of benzoate regulation in the ancestor W3110

In our report of acid evolution, transcriptomes of evolved clones reveal the surprising reversal of acid stress responses (23). We therefore conducted a similar transcriptome analysis of our strains evolved in the presence of benzoate (18). Four of the 2000-generation strains were selected (A1-1, C3-1, G5-1, and G5-2) for comparison with the ancestral strain W3110. RNA was extracted during logarithmic growth from cultures supplemented with a relatively modest benzoate stress (5 mM benzoate) in order to sustain comparable growth of the ancestral strain. In addition, strain W3110 was cultured in medium with no benzoate, in order to identify genes responding to benzoate stress before the generations of benzoate selection.

**Figure 3** plots the log_2_ expression ratios (versus ancestor) for each benzoate-evolved 2000-generation strain against the log_2_ ratios for expression in W3110, with benzoate versus without. Across the genome, genes that were regulated up or down by benzoate in the ancestor showed reversal (by deletion or by down-regulation) in the four independently evolved isolates. In some cases, the reversal involved deletion of several genes or a major regulator, such as the deletion of nearly the entire Gad island *slp*-[*gadW*] in A1-1, or loss of the *gadX* activator in G5-1 and in G5-2. These strains showed the loss or down-regulation of most of the Gad regulon (blue symbols in **Figure 3**). Surprisingly, C3-1 had lost Gad expression, despite the presence of known regulators. Thus, the genotype of C3-1 may reveal novel means of Gad regulation. Overall, reversing the transcriptomic effects of benzoate appeared to be a general property of the evolved strains, and not limited to a few major systems.

**Figure 3.**
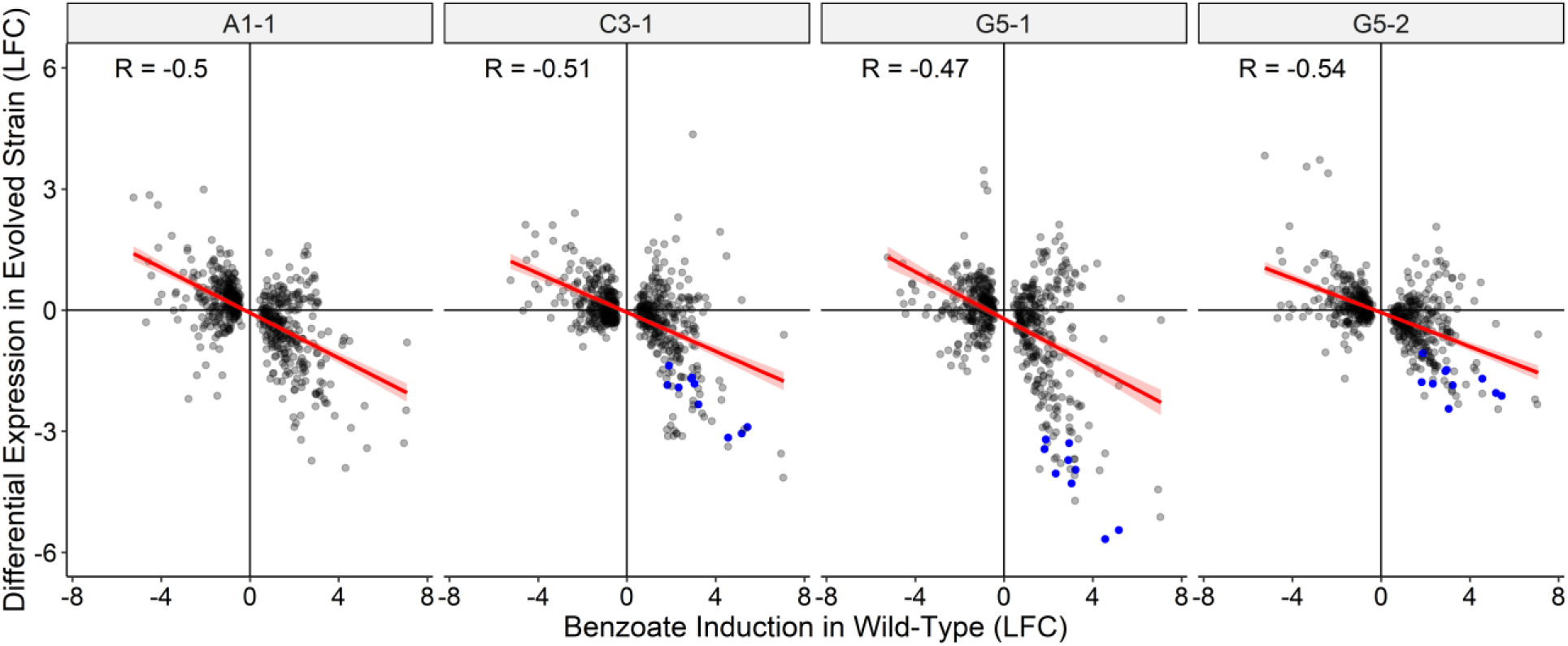
Genes upregulated by benzoate in the ancestral strain W3110 are downregulated in the benzoate-evolved strains. The grey dots indicated the Log_2_ fold changes (LFC), the expression ratios of genes of the benzoate-evolved strains, plotted as a function of LFC for W3110 with or without 5 mM benzoate. Data are from the Supplemental File Table S3. Genes shown are those significantly differentially expressed (p < 0.001) in at least one evolved strain. Genes of the Gad regulon (*gadAEXW, mdtEF, hdeDAB, slp*) and *gadBC* are colored blue. In A1-1 most of the Gad island is deleted (Δ*slp-gadW*). R is the Pearson correlation coefficient.

The log_2_ fold change in expression for individual genes of strain W3110 in the presence or absence of benzoate are presented in **Table S3A**. Genes were deemed significantly differentially expressed if the log_2_ fold change was greater than 1, and p-value less than 0.01 Expression changes are also shown for the 2,000-generation strains versus W3110, all cultured in 5 mM benzoate. Selected differentially expressed genes are shown in **Table 3B-E**.

**Table 3.** Selected differentially expressed genes (Log_2_ fold change ≥ 1, p < 0.01) ^1^

|  | Benzoate /<br>No benzoate<br>W3110 | A1-1 /<br>W3110<br>Benzoate | C3-1 /<br>W3110<br>Benzoate | G5-1 /<br>W3110<br>Benzoate | G5-2 /<br>W3110<br>Benzoate |
| --- | --- | --- | --- | --- | --- |
| Up-<br>regulated | *aceABEFK* | *acs* | *cspB* | *cvpA* | <u>*ompF*</u> |
|  | *appAB* | *glcDE* | *ompG* | *fadB* | *ompL* |
|  | *asr* | <u>*ompF*</u> | *ompL* | *glcDE* | *wcaADK* |
|  | *cyoABCDE* |  | *pspB* | *ompL* | <u>*ymgG*</u> |
|  | *dppCB* |  | *phoE* | *phoE* |  |
|  | *gadABC* |  |  | *phoH* |  |
|  | *gadEF* |  |  |  |  |
|  | *hdeB* |  |  |  |  |
|  | *hyaBF* |  |  |  |  |
|  | *hycABCDEFG* |  |  |  |  |
|  | *marRAB* |  |  |  |  |
|  | *mdtEF* |  |  |  |  |
|  | *narHJ* |  |  |  |  |
|  | *oppABCD* |  |  |  |  |
|  | *slp* |  |  |  |  |
| Down-<br>regulated | *cadAB* | *appBC* | *flu* | *dctR* | *artP* |
|  | *flgBCDEFG* | *dctR* | *gadABC* | *gadABC* | *asr* |
|  | *fliAFN* | *flu* | *hycACDEFGH* | *gadEF* | *flu* |
|  | *fruAK* | *gadABC* | *pepN* | *hdeABD* | *gltIK* |
|  | <u>*ompF*</u> | *gadEF* | *slp* | *hycACDEFGH* | *hycCDEFG* |
|  | <u>*ymgG*</u> | *mdtEF* |  | *mdtEF* | *marRB* |
|  |  | *hycBCDEFGH* |  | *slp* | *slp* |
|  |  | *hdeABD* |  | *sufABCD* | *yahK* |
|  |  | *narHIJ* |  | *yahK* | *yeeR* |
|  |  | *slp* |  | *yhiDM* |  |
|  |  | *yeeR* |  | *wrbA* |  |
|  |  | *yhiDM* |  |  |  |
|  |  | *ymgG* |  |  |  |

<sup>1</sup>All strains were cultured to log phase at pH 6.5, 5 mM sodium benzoate, except for the W3110 control culture without benzoate. Bold letters without underline indicate genes upregulated by benzoate in W3110 but knocked down in benzoate-evolved strains (A1-1, C3-1, G5-1, G5-2). Bold underlined indicates genes downregulated by benzoate in W3110 and that had increased expression in benzoate-evolved strains.

In W3110, benzoate upregulated much of the Gad regulon including both glutamate decarboxylases (*gadA*, *gadB*) and glutamate transporter *gadC*, the regulator *gadE*, and the Gad-associated multidrug efflux pump *mdtEF*. The *hdeABD* portion of the Gad island showed less induction, likely because *hdeAB* is repressed by MarA (**Fig. 1**) which is induced by benzoate (**Table 3**). Benzoate induction of glutamate decarboxylase would be consistent with Foster’s model that cytoplasmic pH depression induces Gad (64). However, the cytoplasmic pH depression caused by benzoate uptake does not involve extreme acidification of the periplasm, since the media are buffered to pH 6.5. Thus, the cell has less need of the periplasmic chaperones HdeA and HdeB.

While MarA is induced by benzoate in the ancestor, the benzoate-evolved strains show loss of various Mar regulon components, either by *marRAB* deletion (strain A5-1, **Table 2**) or by loss of another regulator such as *rob* (strain G5-2). MarA downregulates the large outer membrane porin *ompF* (**Fig. 1**). The downregulation of *ompF* helps exclude toxins and antibiotics (28).

No *ompF* mutations appear in our evolved strains, yet in transcriptomes (**Table 3**) two of the four benzoate-evolved strains (A1-1 and G5-2) showed upregulation of *ompF*. Strain C3-1 upregulated other large porins, *ompG* (65), *phoE* (66), and *ompL* (67). Similarly, strain G5-1 upregulated porins *phoE* and *ompL*. Thus, under long-term exposure to benzoate, with concomitant PMF depletion, all four strains upregulated porins that could enhance fitness by admitting more carbon sources for substrate-level phosphorylation and fermentation. The upregulation of porins could also increase admittance of antibiotics in the benzoate-evolved strains (26, 28).

In strain W3110, benzoate also induced expression of oligopeptide transport operons *dppBC* (68) and *oppABCD* (69). Like the porins, these transporters could increase access to fermentable substrates. These transporters retained expression in the benzoate-evolved strains.

W3110 genes induced by benzoate also included several components of electron transport, including cytochrome *bo3* ubiquinol oxidase (*cyoABCDE*), cytochrome oxidase *bd-II* (*appBC*) (70, 71), nitrate reductase (*narHJ*) (72), and hydrogenase-3 (Hyd-3, *hycABCDEFG*) (73). Hyd-3 converts H+ ions to H2, and associates with the formate-hydrogenlyase complex (FHL); it includes HycA regulator of FHL (74). The acceleration of electron transport is consistent with the effect of uncouplers on respiration, generating a futile cycle during stationary phase (75). The energy loss during stationary phase could explain why strain W3110 enters death phase after several hours in culture with 20 mM benzoate (18).

The FHL/Hyd-3 components (*hyc* genes) were downregulated in all four benzoate-evolved strains. Strain A1-1 had lower expression of *appBC* cytochrome oxidase *bd-II*. All these complexes--Hyd-3, FHL-3, and the *bd-II* oxidase--are generally expressed under low-oxygen conditions (73, 76). Hyd-3 converts 2H^+^ to H_2_, as part of the formate-hydrogen lyase complex which oxidizes formate to CO_2_ (77). Hydrogen production by Hyd-3 is induced by external acid, and is required for extreme-acid survival in low oxygen (78). Thus the benzoate-associated loss of acid-inducible hydrogenase (and of formate-hydrogen lyase) parallels the loss of the acid-inducible Gad system.

No hydrogenases had mutations in our resequenced genomes, although three different mutations appear affecting the nitrate-inducible formate dehydrogenase (*fdnG* and *fdnI*) (79). Also in A1-1, the *narHIJ* quinol-nitrate oxidoreductase (80) is down-regulated (**Table 3; Table S3B**). Thus some kinds of reregulation have occurred that decrease the wasteful expenditure of electrons. These changes most likely enable benzoate adaptation during the low-oxygen period of stationary phase, the period in which the ancestral strain declines (**Fig. 2**).

### Deletion of *slp-gadX, mdtE*, *mdtF*, or *ariR* confers benzoate tolerance

Our resequenced genomes of benzoate-evolved strains offered candidate genes for mechanisms of benzoate tolerance. We selected candidate gene *kanR* replacements from the Keio collection to run batch growth curves that approximate the daily experience of serial dilution. Growth curves can reveal fitness differences of several types: duration of lag phase, rate of log-phase growth, end-point reached at stationary phase, and post-stationary death rate.

We started with the isolates from population A1, which suggests a progression of acquired mutations that contribute to benzoate tolerance (**Table S2**). The early-generation isolates do not necessarily represent direct ancestors of the later strains, yet the *slp-gadX* deletion was present in all A1 strains sequenced, and this region encompasses most early mutations found in other populations (**Table 2**). We constructed, by recombineering (81), a strain with a deletion spanning the start of *slp* to the end of *gadX* (*slp-gadX* strain).

Strains were cultured in microtiter plates with 15 mM benzoate, as described under Methods (**Fig. 4**). The OD_600_ value was measured at 16 h to evaluate the endpoint; and for each condition, we show a replicate curve with median 16-h OD_600_. All eight replicate curves are shown in **Fig. S2**. The *slp-gadX* strain had higher 16-h growth relative to W3110, and nearly as high as that of the benzoate-evolved strain A1-1 (**Fig. 4A**). W3110 mutants deleted for *gadE* or *gadX* did not show a consistent effect on growth, although their regulation of the Gad island (64) might have fitness effects detectable by direct competition.

**Figure 4.**
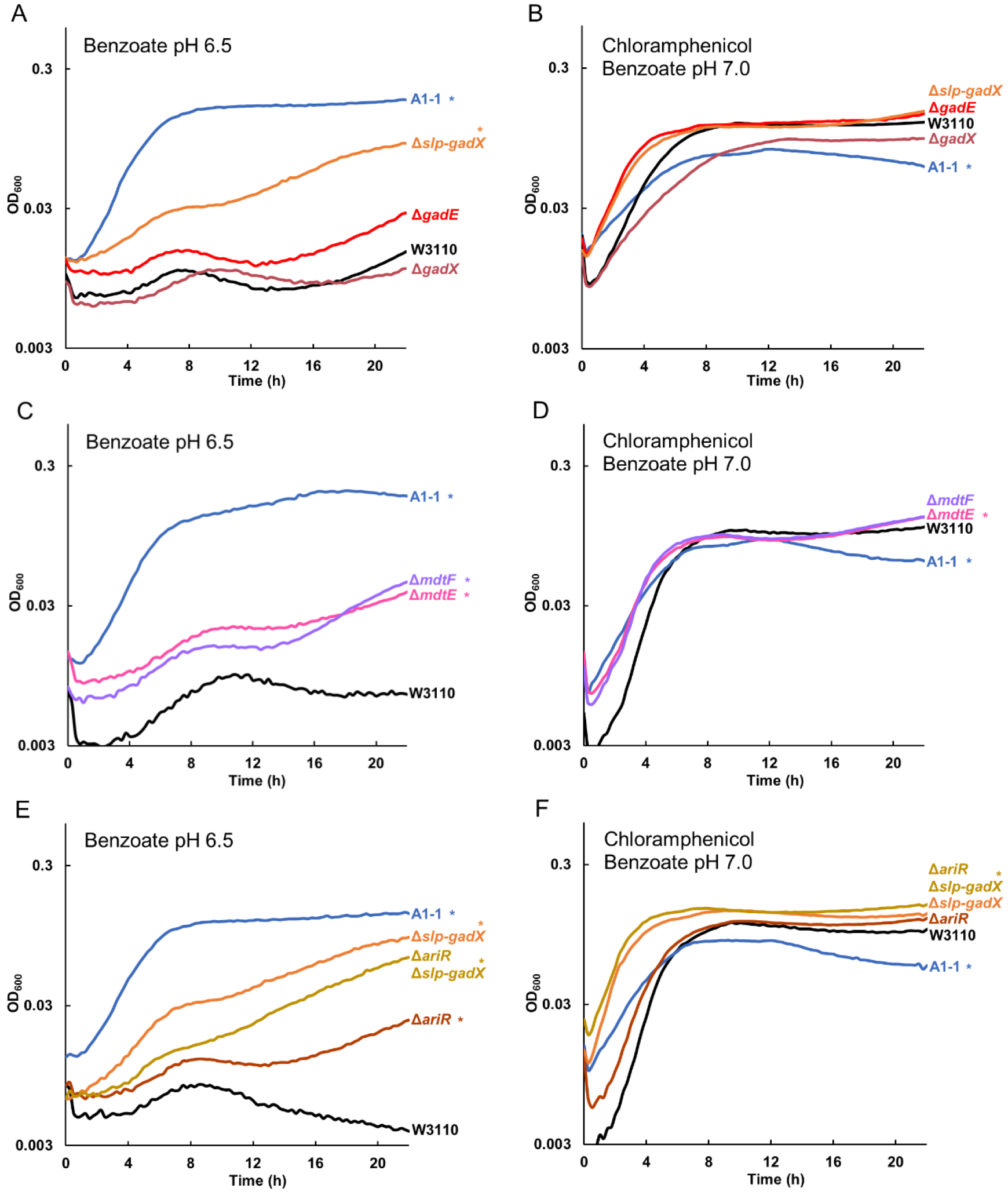
Gad regulon and related mutations in strain A1-1 increase growth in benzoate. Replicate curves are shown in **Fig. S4**. Strains were cultured overnight in LBK 100 mM PIPES pH 6.5 (A, C, E) or LBK 100 mM MOPS pH 7.0 (B, D, F), both supplemented with 5 mM benzoate. Eight replicate samples from two overnight cultures (four from each) were diluted 1:200 in a 96-well plate into fresh medium; LBK 100 mM PIPES pH 6.5 supplemented with 15 mM benzoate (A, C, E) or LBK 100 mM MOPS pH 7.0 supplemented with 5 mM benzoate and added 4 μg/mL chloramphenicol (B, D, F). For each group of replicates, a curve with median OD_600_ at 16 h is presented. Asterisk indicates significant difference from W3110 at 16 h, Tukey’s test p ≤ 0.05, n=8. All eight replicates are shown in **Figure S4. A.** W3110 constructs Δ*slp-gadX*, Δ*gadE::kanR*, and *ΔgadX::kanR* were cultured in 15 mM benzoate alongside parent strain W3110. The Δ*slp-gadX* strain grew to a higher OD_600_ than did W3110 at 16 h, though not as high as benzoate-evolved strain A1-1. **B.** Strain A1-1 and W3110 constructs Δ*slp-gadX*, Δ*gadE::kanR*, and Δ*gadX::kanR* were cultured with W3110 in benzoate and chloramphenicol. Strain A1-1 grew significantly less than W3110 at 16 h. **C.** *ΔmdtE::kanR* and Δ*mdtF::kanR* strains outgrew W3110 in benzoate, but did not grow as high as A1-1. **D.** Δ*mdtE::kanR* and Δ*mdtF::kanR* strains grew higher than W3110 in 5 mM benzoate with chloramphenicol, whereas strain A1-1 reached a lower OD_600_ at 16 h. **E.** Δ*ariR::kanR* had increased growth in benzoate, but not as high as Δ*slp-gadX*, Δ*ariR::kanR* Δ*slp-gadX*, or A1-1. **F.** While Δ*ariR::kanR* and Δ*slp-gadX* did not affect growth in benzoate with chloramphenicol, Δ*ariR::kanR* Δ*slp-gadX* outgrew the ancestor W3110.

Salicylate has a similar effect to benzoate on the growth of W3110 and the benzoate-evolved strains, and its effect starts at a lower concentration (18). An experiment similar to that of **Figure 4B** was conducted using chloramphenicol with 2 mM sodium salicylate instead of 5 mM benzoate; similar results were obtained. **Figure S3** presents salicylate experiments comparable to all the chloramphenicol-benzoate experiments in Figures 4, 5, 6, 7.

**Figure 5.**
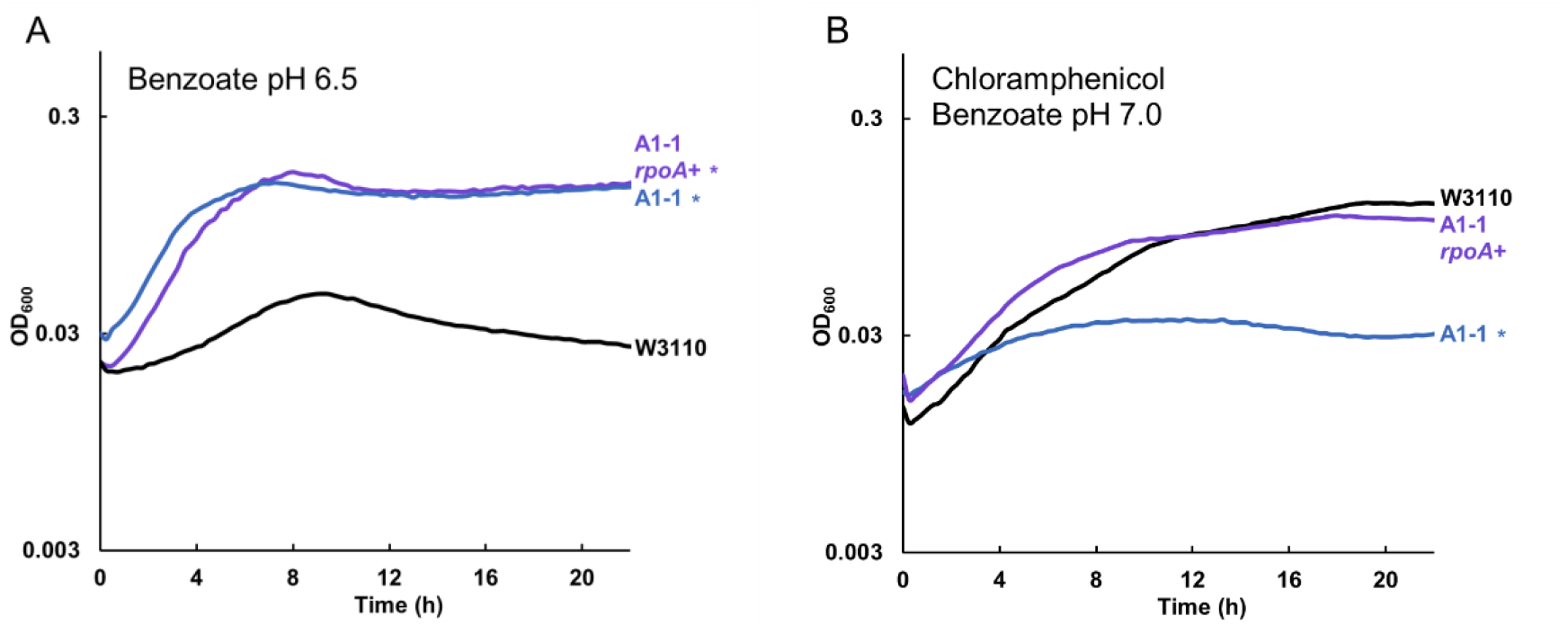
Chloramphenicol sensitivity with *rpoA* K271Q replacement by *rpoA*^+^ in A1-1. Replicate curves are shown in **Figure S5. A.** Construct A1-1 with *rpoA* K271Q reversion to *rpoA*^+^ (A1-1 Δ*yhdN::frt rpoA*^+^) showed no significant difference from A1-1 (cultured with 15 mM benzoate). **B.** A1-1 Δ*yhdN::frt rpoA*^+^ grew to a comparable level as W3110 in 8 μg/mL chloramphenicol with 5 mM benzoate. Culture conditions for panels A, B were the same as for Figure 4 A, B respectively, except that the chloramphenicol concentration was 8 μg/mL (B). Significant difference from W3110 was determined at 16 h, Tukey’s test p ≤ 0.05, n=8.

We then tested the benzoate tolerance of strains deleted for the *mdtE* and *mdtF* components of the MdtEF-TolC MDR efflux pump, which is included in the acid fitness island deletion (29). Knockout strains *mdtE::kanR* and *mdtF::kanR* each reached a higher 16-h density relative to W3110 in 15 mM benzoate at pH 6.5 (**Fig. 4C**). However the increase was significantly less than that found for strain A1-1 and the *slp-gadX* strain. This suggests that various genes of the Gad island make additive partial contributions to benzoate tolerance.

Another mutation in A1-1 that had the potential to affect benzoate tolerance was a transposon-mediated knockout of *ariR* (18, 43). AriR (YmgB) regulates both Gad acid resistance and biofilm formation (43) possibly mediated by RpoS (82). A knockout strain *ariR::kanR* had increased endpoint growth compared to wild-type. We then tested whether an *ariR* deletion interacted with the acid fitness island deletion in an additive manner, or whether its function was made redundant by the the *slp-gadX* deletion. We transduced *ariR::kanR* into the *slp-gadX* strain (**Fig. 4E**). This strain grew identically to the *slp-gadX* strain in 15 mM benzoate, suggesting the *ariR* single deletion increases benzoate tolerance through regulation of the Gad island.

### Chloramphenicol sensitivity is conferred by *rpoA* K271Q but not by *slp-gadX* deletion

Benzoate tolerance of our benzoate-evolved strains is associated with a tradeoff of sensitivity to chloramphenicol, as seen for strains A1-1, A5-1, C3-1, G5-1, and G5-2 (18). We sought to clarify how benzoate tolerance is connected with antibiotic sensitivity. Our new isolates from the early generations confirmed that chloramphenicol sensitivity emerged later than benzoate tolerance; only one generation 1,400 isolate (A1-5) and none of the earlier isolates showed measurable decrease in growth with chloramphenicol relative to W3110, at 4 μg/ml or at 8 μg/ml (**Table 2**, second row). The *slp-gadX* deletion and deletions of individual genes were also tested for their effects on chloramphenicol resistance (**Fig. 4B, D, F**). Chloramphenicol resistance was tested in a medium containing a low level of benzoate (5 mM) in order to induce any Mar and non-Mar MDR systems. None of the mutations affected chloramphenicol resistance, despite the known chloramphenicol efflux by MdtEF-TolC (52). This could occur because chloramphenicol is exported by redundant systems outside *slp-gad*, including AcrAB-TolC.

For strain A1-1, we hypothesized that its chloramphenicol sensitivity required an additional mutation acquired relatively late in the evolution experiment, perhaps between generations 900 and 1,400. An interesting candidate was the *rpoA* (RNA-polymerase alpha subunit) mutation that appeared in strain A1-5 (generation 1,400, chloramphenicol sensitive) and persisted in A1-1. The *rpoA* gene encodes the α-subunit of RNA polymerase. The carboxy-terminal domain of this subunit regulates gene expression by interacting with upstream promoter elements, and transcription factors such as CRP.

Our *rpoA* mutation K271Q is nearly the same as that of a well-studied *rpoA* allele, *rpoA341* K271E (83, 84). The *rpoA341* allele downregulates certain positively-controlled regulons, such as *mel, ara*, and cys; but not all positive regulators are affected. The *rpoA341* allele may also upregulate loci such as *ompF* (83). We found *ompF* upregulated in strain A1-1 (**Table 3, Table S3B**). Strain A1-1 also had decreased expression of *araBAD* and of *cysA*, as reported for the *rpoA341* allele (83).

We sought to test the contribution of *rpoA* K271Q in strain A1-1 with respect to benzoate tolerance and chloramphenicol sensitivity. To do this, we replaced the mutant allele with *rpoA^+^* by co-transduction from W3110 into A1-1 using the linked marker Δ*yhdN::kanR*. The *yhdN::kanR* marker had no effect on growth of strain A1-1, with or without chloramphenicol (**Fig. S4**). Strain JLS1616 (A1-1 Δ*yhdN rpoA*^+^) was cultured in parallel with strain A1-1 and the parent W3110 (**Fig. 5**; replicate curves are shown in **Fig. S5**). In 15 mM benzoate, the A1-1 *rpoA*^+^ strain grew similarly to the parent A1-1, with no significant difference at any time point (panel A). However, with chloramphenicol and low benzoate (5 mM) the A1-1 *rpoA*^+^ showed growth comparable to that of W3110, whereas A1-1 with the original *rpoA* mutation was sensitive to chloramphenicol. Thus, *rpoA* K271Q is responsible for the loss of chloramphenicol resistance associated with benzoate evolution of A1-1.

### *cspC* deletion confers benzoate tolerance, and *rfaY* deletion decreases growth rate in chloramphenicol

The C3-1 strain has two transposon-mediated deletions that might affect growth in benzoate, affecting a post-transcriptional regulator *cspC* and the LPS kinase *rfaY* (18). The sequence of *cspC* is similar to that of other cold-shock proteins, and the CspC protein has been shown to upregulate genes under RpoS control by stabilizing the *rpoS* mRNA (85–87).

A Δ*cspC::kanR* knockout strain showed increased growth relative to strain W3110 in 15 mM benzoate (**Fig. 6A**; replicate curves, **Fig. S6**). However, in media with 4 μg/mL chloramphenicol, there was no significant difference in growth (**Fig. 6B**). Thus, we found evidence for a contribution of Δ*cspC* to benzoate tolerance of C3-1 but not to chloramphenicol sensitivity.

**Figure 6.**
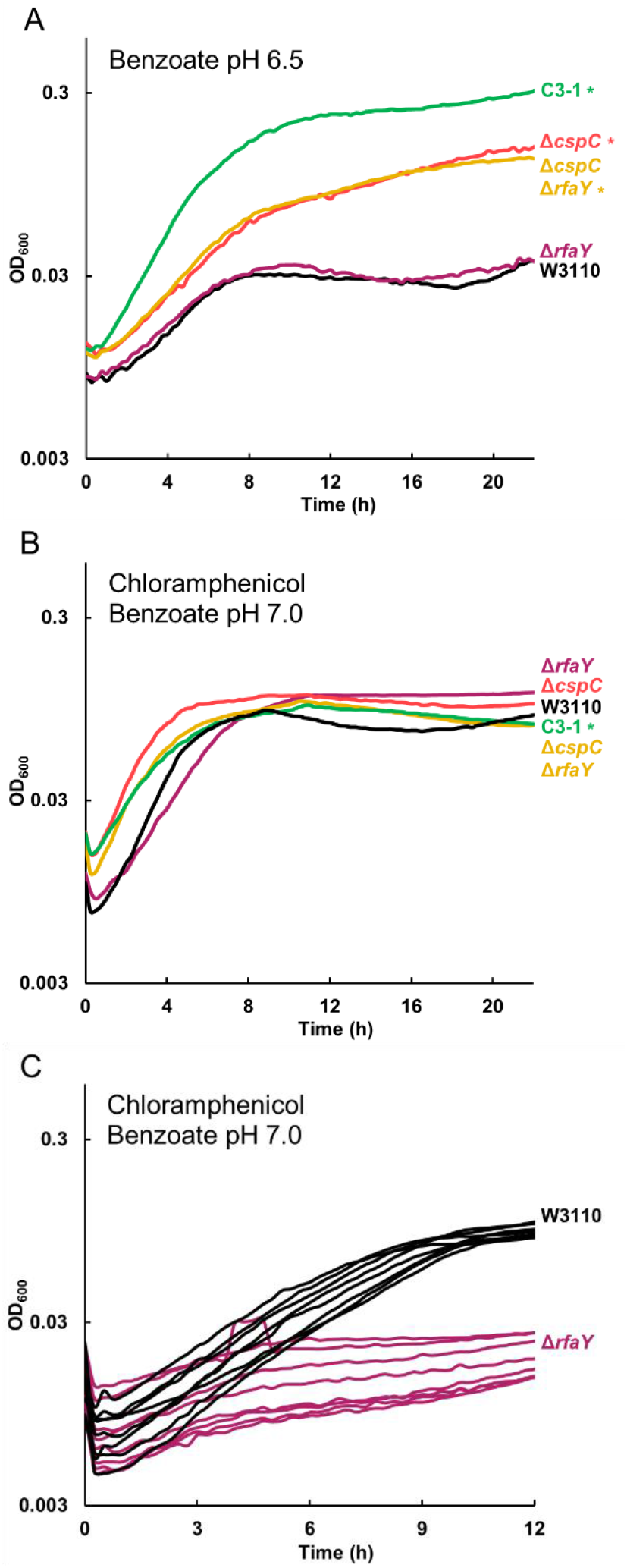
*ΔcspC* deletion confers benzoate tolerance and *ΔrfaY* confers chloramphenicol sensitivity. Replicates are shown in **Figure S6. A.** In 15 mM benzoate, W3110 Δ*cspC* reached higher growth than W3110, but W3110 Δ*rfaY* showed no growth difference from W3110. The Δ*cspC* Δ*rfaY* growth curve resembled that of Δ*cspC*. **B.** W3110 constructs Δ*rfaY::kanR* and Δ*cspC:.kanR* Δ*rfaY.:frt* showed no difference from W3110 at 16 h, but Δ*rfaY::kanR* had a lower log-phase growth-rate compared to W3110 in 5 mM benzoate with 4 μg/mL chloramphenicol. **C.** At 8 μg/mL chloramphenicol, 5 mM benzoate, W3110 Δ*rfaY::kanR* has lower log-phase growth rate than W3110 (t-test p < 0.01); for each strain, 8 individual replicates are shown. Significant difference from W3110 was determined at 16 h, Tukey’s test p ≤ 0.05, n=8.

An Δ*rfaY::kanR* strain showed no phenotype in 15 mM benzoate (**Fig. 6A**). The double mutant W3110 Δ*cspC::kanR ΔrfaY::frt* grew identically to W3110 Δ*cspC*. In chloramphenicol however, W3110 Δ*rfaY::kanR* had a mean growth rate of 0.12 ± 0.01 doublings per hour in early log phase, which was less than the mean W3110 growth rate of 0.39 ± 0.02 doublings per hour (**Fig. 6C**). A t-test comparison gave a p-value of p < 0.01. The double mutant also showed a significant decrease of growth rate in chloramphenicol, compared to the single mutant Δ*cspC::kanR*. Thus, the double mutant showed the *cspC* phenotype cultured in 15 mM benzoate but the Δ*rfaY* phenotype cultured with chloramphenicol (5 mM benzoate).

The C3 population isolates has no mutations within *slp-gadX*, except for a 76-bp deletion of *gadX* found in C3-5. Yet, the C3-1 transcriptome showed lower expression of *gadABC* (**Fig. 3**) so its genome must have altered regulation of acid-tolerance pathways. Given that *cspC* is implicated in the stabilization of the *rpoS* mRNA (86, 88), and RpoS regulates Gad (**Fig. 1**) it is possible that *cspC* deletion downregulates Gad via decreased RpoS concentration.

### *emrA* and *add* deletions confer benzoate tolerance, while *rob* and *hfq* confer chloramphenicol sensitivity

From population G5 (**Table 2**), strains G5-5 and G5-1 have a knockout mutation in *emrA*, which normally encodes part of the EmrAB-TolC pump (19). The *emrA* gene acquires point mutations--but not knockouts--under evolution with the uncoupler CCCP, which is expelled by EmrAB-TolC (20). We found that deletion of *emrA* conferred a degree of tolerance to benzoate (**Fig. 7A**; replicate curves, **Fig. S7**). The *emrA* deletion confers added benzoate tolerance even in the presence of chloramphenicol (**Fig. 7B**).

**Figure 7.**
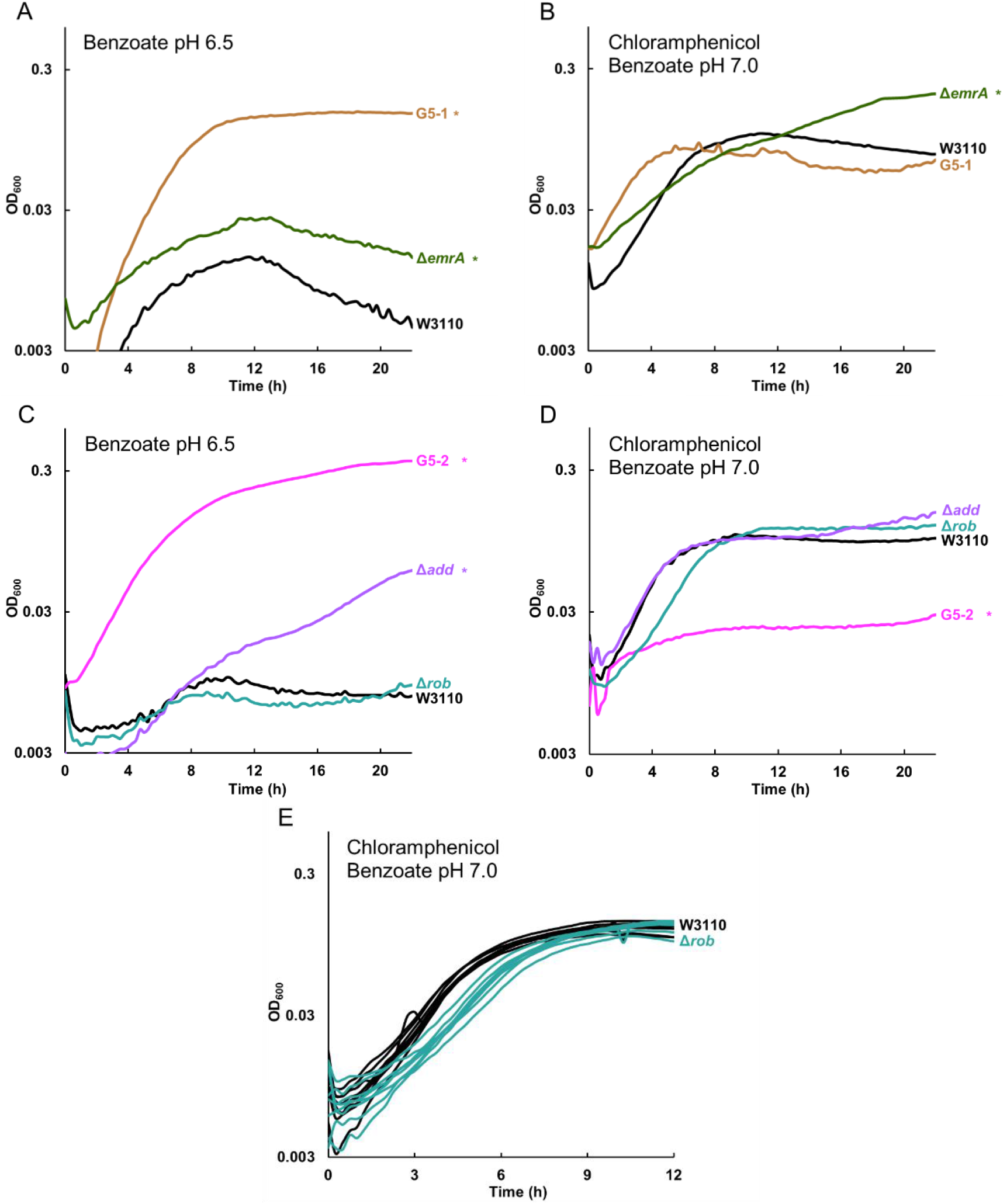
*Δadd* confers benzoate tolerance, and Δ*rob* decreases log-phase growth rate in chloramphenicol. Replicates are shown in **Figure S7. A.** Strains G5-1 and W3110 Δ*emrA::kanR* outgrew W3110 at 16 h, in 15 mM benzoate. **B.** W3110 Δ*emrA* outgrew W3110 and G5-1 at 16 h, in 5 mM benzoate, 4 μg/ml chloramphenicol. **C.** G5-2 and W3110 Δ*add::kanR* outgrew W3110 in 15 mM benzoate. Δ*rob::kanR* did not affect growth. **D.** W3110 Δ*add::kanR*, W3110 Δ*rob::kanR*, and W3110 showed no difference in 5 mM benzoate, 4 μg/mL chloramphenicol. Strain G5-2 grew less than W3110. **E.** Δ*rob::kanR* conferred a lower log-phase growth rate in 5 mM benzoate, 4 μg/mL chloramphenicol (t-test p < 0.01). 8 replicates per strain are shown.

Another strain from G5 population, G5-2, shows the greatest sensitivity to chloramphenicol and tetracycline, of all 2,000-generation strains tested (18). G5-2 contains a frameshift in *add*, a gene encoding adenosine deaminase which catalyzes a proton-consuming reaction of purine catabolism that protects the cell from acid by a mechanism analogous to that of GadA (89). Deletion of *add* conferred partial tolerance to benzoate (**Fig. 7C**). This result is yet another example of benzoate selection against an external-acid protection mechanism. G5-2 also contains a point mutation in *rob* encoding a MarA-type regulator that enhances chloramphenicol resistance (45). Deletion of *rob* did not significantly enhance growth in 15 mM benzoate (**Fig. 7C**) but did decrease the log-phase growth rate in chloramphenicol, from an average W3110 growth rate of 0.63 ± 0.01 doublings per hour to an average growth rate of 0.59 ± 0.02 doublings per hour for W3110 *Δrob::kanR* (**Fig. 7E**). A t-test comparison gave a p-value of p < 0.01. Replicates for the log-phage growth region are shown for W3110 *Δrob::kanR* and W3110 in 4 μg/ml chloramphenicol (**Fig. 7E**).

G5-2 also contains a missense mutation in *hfq*, which encodes a pleiotropic regulator that functions as an RNA chaperone (90) and DNA-binding protein (91). Hfq is associated with antibiotic resistance and is a target for antimicrobial chemotherapy (92, 93). We found that an *hfq* deletion confers partial tolerance to 15 mM benzoate (**Fig. 8A**) and sensitivity to chloramphenicol (**Fig. 8B**). In chloramphenicol, the growth curve of W3110 *Δhfq::kanR* was indistinguishable from that of G5-2. Thus, *hfq* mutation could contribute a major part of the G5-2 phenotype of benzoate tolerance associated with antibiotic sensitivity.

**Figure 8.**
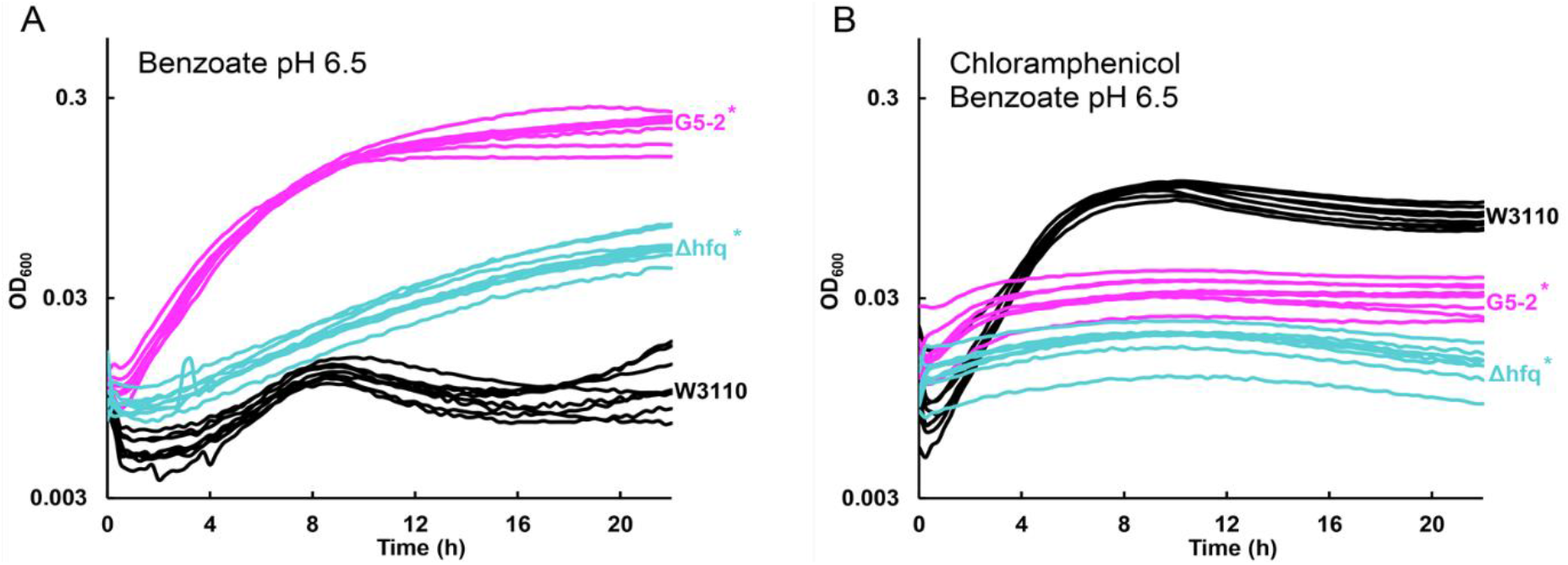
*Δhfq::kanR* increases growth in 15 mM benzoate and decreases growth in chloramphenicol. **A.** Strains G5-2 and W3110 Δ*hfq::kanR* outgrew W3110 at 16 h, in 15 mM benzoate. **B.** W3110 outgrew strains G5-2 and W3110 *Δhfq::kanR* at 16 h, in 5 mM benzoate, 4 μg/ml chloramphenicol.

### Hydrogenase 3 deletion enhances late growth

Our transcriptomes showed that benzoate induces hydrogenase 3 in W3110, but that all four benzoate-evolved strains lose expression of *hycEFG* (**Table 3**). We tested the growth of W3110 strains deleted for *hycE* (encoding the hydrogenase activity, 2H^+^ → H_2_) and for *hycF* (subunit of formate hydrogenlyase complex) (94). Both *ΔhycE::kanR* and *ΔhycF::kanR* strains grew similarly to W3110 until stationary phase. At 16 hr, there was no significant difference between the ancestral and mutant OD_600_. However, after 20 hr, when hydrogenase would be active, the mutants achieved a higher OD_600_ than did W3110 (**Fig. 9**). These results confirm that although benzoate exposure induces consumption of cytoplasmic protons via hydrogenase 3, as part of formate breakdown, this activity decreases long-term relative fitness in benzoate. By contrast, the hydrogenase 1 (*hya*) consumes H_2_, producing 2H^+^ (73).

**Figure 9.**
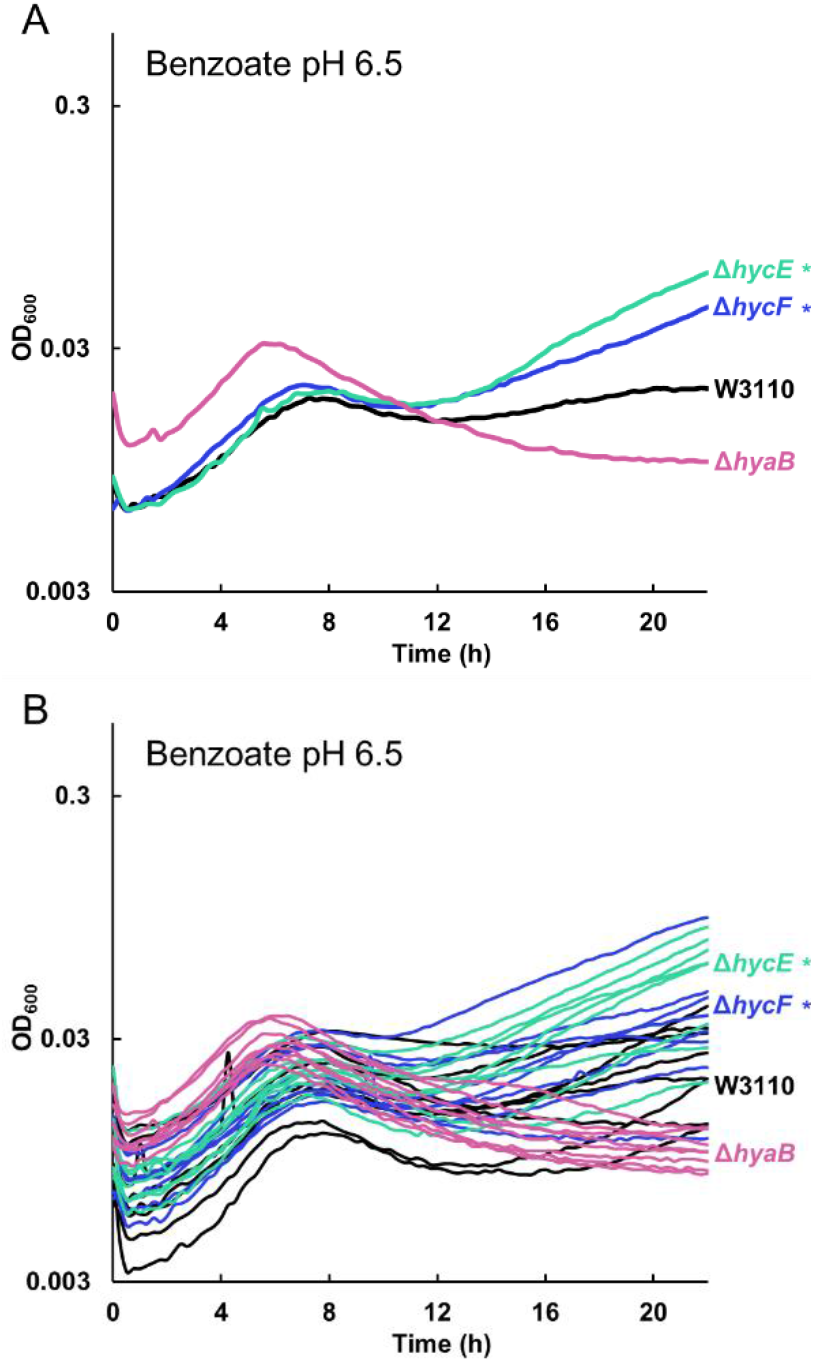
Deletion of hydrogenase Hyd-3 (*hyc*) enhances late growth in 15 mM benzoate pH 6.5. W3110 with Δ*hycE::kanR* or Δ*hycF::kanR* grows to higher median value of OD_600_ at 20 h than that of the parent W3110. The W3110 Δ*hyaB::kanR* strain deleted for Hyd-1 shows no enhancement of late growth compared to W3110. **A.** Median growth curves presented for each set of 8 replicates. **B.** All 8 replicates shown for each strain.

### Other genes tested

KanR knockout strains were tested for other genes that had mutant alleles in our benzoate-evolved isolates, but no significant difference from the ancestor was detected for growth with benzoate or with chloramphenicol. As noted, such differences might emerge under extended direct competition. The genes tested by 22 h culture with 15 mM benzoate include: *acnA, chbC, cpxA, deaD, fucA, mdtA, gltP, hdeD, pepN, rnb, yhfM, uxaA*, and *yhiD*.

## DISCUSSION

Here we combined genomic, transcriptomic, and genetic approaches to study how *E. coli* evolve in the presence of benzoate and salicylate. Our work extends the picture of known MDR-related genes mutated under exposure to partial or full uncouplers (18)(20) and suggests additional mechanisms for the increased sensitivity to antibiotics, such as the upregulation of porins and pleiotropic regulators. This subject has implications beyond *E. coli*, because the microbiomes of soil and rhizosphere (95) as well as human and animal digestive tracts (96) are exposed to benzoate and related molecules.

### Evolution in the presence of benzoate reverses short-term benzoate stress response

We confirmed by analysis of candidate gene deletions that despite short-term benzoate upregulation of Gad genes (**Table 3**) the presence of benzoate actually selects against the Gad island (*slp-gadX*) and against Gad components *mdtE* and *mdtF*, which encode a multidrug efflux pump (**Fig. 4**). Our transcriptomes reveal a striking pattern of repression of Gad as well as other benzoate-inducible gene products in the benzoate-evolved strains (**Table 3**). Even a strain with no mutations in the fitness island (C3-1) had downregulated Gad, possibly by mutation of the RpoS post-transcriptional activator CspC. Similarly, strain G5-2 could have downregulated Gad via deletion of *rob* or of *hfq* (which activates RpoS). Collectively, these results show a pattern of convergent evolution achieved via one of several possible genetic mechanisms.

Our transcriptomes also showed that evolution with benzoate <u>increased</u> the expression of several large-hole porins (OmpF, OmpG, PhoE, OmpL). These porins amplify access to fermentable carbon sources for substrate-level phosphorylation, thus decreasing dependence on PMF. But OmpF is normally downregulated by benzoate derivatives via MarA, in order to exclude antibiotics (26). It is interesting that benzoate induced short peptide transporters *opp* and *dpp* (**Table 3**) which offer another means of access to fermentable carbon sources; this expression of peptide transporters was maintained in benzoate-evolved strains. For *opp*, there is controversial evidence that increased peptide transport coincides with antibiotic sensitivity (97–99).

Another form of reversal of benzoate regulation was the downregulation of hydrogenase 3 (**Table 3, Fig. 9**). The Hyd-3 substrate, formate, arises from *E. coli* fermentation, increasing in stationary phase. Hyd-3 as part of formate hydrogenlyase complex normally converts formate to CO2 and H2. Yet we show that in the presence of benzoate, Hyd-3 deletion enhances fitness during late stationary phase, the period where we would expect FHL/Hyd-3 to be active (76). There is evidence that FHL activity exports protons (100). If so, this could be yet another system that wastes energy as exported protons drive more benzoate into the cell—and thus decreases relative fitness over long-term subculturing. The possible effects of benzoate and other partial uncouplers could be relevant to the biotechnology of hydrogen production (94).

In strain A1-1, several alternative terminal oxidases were downregulated. Antibiotic resistance is linked to bacterial respiration, as several bactericidal antibiotics are shown to increase respiratory rates, while most bacteriostatic compounds decrease respiration (101). One means of adaptation to high benzoate concentration could be to limit respiration by down-regulating unneeded components of electron transport, including anaerobic respiration. This would limit the energy wasted by uncoupling respiration from ATP synthase. Introduction of antibiotics to a respiration-compromised cell could amplify the phenotype, and lead to benzoate-induced antibiotic sensitivity.

Reversal of short-term stress response following long-term serial culture has been seen in other evolution experiments as a tendency to restore global pre-stress conditions. After heat-stress evolution of *E. coli*, resulting strains show loss of heat shock gene expression (102). The fitness cost of stress response could involve either the cost of stress-induced gene expression, when it fails to provide a benefit (22); or the cost of excess PMF expenditure by a transporter or an efflux pump (25).

### Growth of candidate-gene deletant strains reveals contributions to benzoate tolerance and chloramphenicol sensitivity

The loss of MdtEF-TolC drug efflux system directly increases benzoate tolerance. This efflux pump couples drug efflux to PMF (31, 33, 103). Theoretically, benzoate decreases PMF by shuttling protons through the membrane. Thus, the presence of other proteins that utilize PMF further depletes the pool of extracellular protons available for core cell processes. It is possible that deleting drug efflux systems decreases proton flux through systems depleted by benzoate; thus the fitness cost of drug pumps is amplified by benzoate.

We showed four candidate alleles associated with antibiotic sensitivity, three of which have global or pleiotropic effects: *rpoA* mutation in A1-1; the *rob* mutation in G5-2; and *hfq*,whose deletion conferred both tolerance to benzoate and sensitivity to chloramphenicol. Hfq deletion is a known source of drug sensitivity (92, 93).

Since the mutation in *rpoA* is in the carboxy-terminal domain, which interacts with UP-elements and certain transcription factors, it is possible that this allele downregulates a large set of genes (104, 105). This downregulation could free up resources for response to benzoate stress, which could include a large number of PMF-driven antibiotic resistance genes. In fact, there is previous evidence suggesting that an interaction between the CTD of RpoA and MarA is necessary for MarA to induce the Mar regulon (105). Mutation of the RpoA CTD could block MarA activation of drug resistance despite *marA* transcription.

The *rfaY* gene (which is mutated in strain C3-1) encodes a membrane-bound enzyme that phosphorylates the inner core of lipopolysaccharide (LPS), a function that has been implicated in membrane stability (106). Decreased membrane stability caused by the *rfaY* mutation may increase the permeability of certain antibiotics, thereby decreasing antibiotic resistance of strain C3-1.

Another surprising finding was the pervasive occurrence of small mutations in genes for aromatic biosynthesis and catabolism, such as a point mutation were found in *folD* and 1 base-pair deletion was detected in *add*. The *folD* gene is essential (53) and could not be deleted, but deletion of *add* was shown to enhance benzoate tolerance. The evidence points to further exploration of the role of benzoate and salicylate in modulating efflux of aromatic intermediates of metabolism (62) especially given the benzoate-evolved enhancement of substrate influx via porins (**Table 3**).

Note that the relative fitness advantage of a given allele can accrue by various means at different phases of the growth cycle. For most of the candidate genes we tested, such as *gadE, mdtE*, and *cspC*, deletion enabled cells to grow to a higher optical density than the parent W3110. However, the chloramphenicol sensitivity associated with some alleles, such as *rfaY*, was caused by slower rate of growth during log phase. While the bacteria grew to an optical density comparable to that of W3110, had the two strains been competing in co-culture, the mutant strain would have soon lost out to the parent.

Overall, we reveal genetic mechanisms by which multigenerational exposure to benzoate leads to increased tolerance of benzoate or salicylate, with the tradeoff of sensitivity to certain antibiotics. Our findings have implications for the roles of benzoate as a food preservative, and for salicylate as a plant defense signal and as a therapeutic agent.

## METHODS

### Bacterial strains and culture conditions

*E. coli* K12 W3110 (Laboratory stock D13) was the parent for all genetic analysis. The strain was resequenced for analysis (18). Unless otherwise specified, bacteria were cultured in LBK (10 g/L tryptone, 5 g/L yeast extract, 7.45 g/L KCL) with a pH buffer, at 37 °C. Growth media were supplemented with benzoate, salicylate, kanamycin (50 μg/ml), or chloramphenicol (4 or 8 μg/ml) as necessary. For growth curves, media was buffered to pH 7.0 with 100 mM (3-(N-morpholino)propanesulfonic acid) (MOPS; pKa = 7.01) or to pH 6.5 with 100 mM piperazine-N,N’-bis(2-ethanesulfonic acid) (PIPES; pKa = 6.66), containing 70 mM Na+. The media pH was adjusted with either 5 M HCL or 5 M KOH. Cultures were incubated at 37°C unless otherwise specified. Strains with *kanR* insertions were obtained from the Keio collection (107). The XTL241 strain containing the *cat-sacB* fusion was obtained from the Court lab at NCI (81). All other strains used in this study are derived from W3110 and listed in **Table 1**, and in **Table S1** (new isolates with resequenced genomes).

### Growth Curves

For growth curves in 15 mM benzoate, strains were cultured overnight in LBK 100 mM PIPES pH 6.5 supplemented with 5 mM benzoate. These cultures were diluted 1:200 in a 96-well plate into fresh LBK buffered to pH 6.5 with 100 mM PIPES supplemented with 15 mM benzoate. OD_600_ was read in a Spectramax spectrophotometer every 15 min for 22 h. For growth curves in chloramphenicol, strains were cultured overnight in LBK 100 mM MOPS pH 7 supplemented with 5 mM benzoate as needed. These cultures were diluted 1:200 into fresh media supplemented with 4 μg/mL chloramphenicol, unless stated otherwise. Endpoint OD_600_ was defined as the cell density after 16 h. Significance tests included ANOVA with Tukey’s post-hoc test (R software). Each data figure represents three experiments in each of which 8 replicate cultures were tested, unless stated otherwise.

### Genome sequencing of early generation evolved-strains

Genomic DNA from early-generation clones and from the ancestral stock W3110 was extracted with a DNeasy DNA extraction kit (Qiagen) and a MasterPure Complete DNA and RNA purification kit (Epicentre, WI). Purity was determined by Nanodrop 2000 spectrophotometer (Thermo Fisher Scientific), and concentration was determined by Qubit 3.0 fluorometer (Thermo Fisher Scientific).

Genomic DNA was sequenced on an Illumina MiSeq platform, at the Michigan State University Research Technology Support Facility Genomics Core. Libraries were prepared with an Illumina TruSeq Nano DNA library preparation kit. After library validation and quantitation, libraries were pooled and loaded on an Illumina MiSeq flow cell. Sequencing was performed in a 2 by 250-bp paired-end format with an Illumina 500 cycle V2 reagent cartridge. Base calling was performed by Illumina Real Time Analysis v1.18.54, and the output of RTA was demultiplexed and converved to FastQ format with Illumina Bcl2fastq v1.8.4. Mutations were called by alignment to the *E. coli* W3110 reference NC_007779.1 using the *breseq* computational pipeline (49).

### Strain construction

*E. coli* W3110 strains were constructed by P1 transduction and by recombineering. P1 transduction was performed by standard methods (108). Strains carrying *kanR* resistance cassettes in genes of interest were acquired from the Keio collection (107). Insertions were verified by PCR-amplification of the interface between the kanR allele and the area surrounding the insertion.

The *slp-gadX* strain (JLS1732) was constructed using lambda-red recombineering according to the protocol by Thomason (81). Generation of the acid fitness island knockout strain, JLS1732, was performed using λ recombineering with the protocol described by Thomason *et al*, 2014. Overnight cultures of E. coli with the pSIM6::ampR plasmid were diluted 1 to 70 in LB (5 g/L NaCl) and grown to mid log-phase (OD_600_ between 0.4 and 0.6) in a shaker flask at 32°C. A 15 mL aliquot of this subculture was transferred to a fresh flask and shaken at 42 °C for 15 min. Cells were made electro-competent and electroporated with a DNA oligonucleotide. Cells were outgrown at 32°C for 3 to 5 hours, and plated onto selective media. For construction of JLS1732 (Δ*slp*-Δ*gadX*), a dsDNA oligo containing a cat-sacB (chloramphenicol resistance – sucrose sensitivity selection/counter-selection marker) with 50 bp of homology to *slp* and 50 bp of homology to *gadX* was constructed using the *cat-sacB* hybrid primers listed in **Table 2**. Then, this region was replaced with the 70-bp oligo, aaacagtaatatgtttatgtaatattaagtcaactaatagatatttctttatagttttcatctgattctg, to produce a strain with a clean break at the start of *slp* and end of *gadX*.

### Transcriptome analysis

The transcriptomes of evolved isolates in comparison with ancestor were obtained as for Ref (23). For RNA extraction, bacteria were cultured to stationary phase in LBK buffered to pH 6.5 with 100 mM PIPES at 37°C. Cultures were diluted 1:50 into fresh medium supplemented with 5 mM potassium benzoate, and grown to early log phase (determined by OD_600_ of 0.4). At mid-log phase (OD_600_ between 0.4 and 0.6), cultures were diluted 6:1 into 5% phenol-ethanol solution and pelleted. The pellet was resuspended in TE buffer (100 μL) with 3 mg/mL lysozyme, as described by He *et al* (23). A Qiagen RNeasy minikit was used to further purify RNA. An additional DNase treatment (MoBio DNase-Max) was conducted.

Illumina RNA-Seq libraries were constructed for sequencing. An enrichment of messenger RNA was achieved by depleting ribosomal RNA (rRNA) by following the guidelines of the Ribo-Zero rRNA Removal Kit (Illumina)(23, 109). The RNA-Seq library was prepared via the ScriptSeq v2 RNA-Seq library preparation kit (Epicenter, WI) with a starting concentration of 15 ng rRNA depleted RNA for each library. The resulting random-primed cDNA was purified with the MinElute PCR Purification Kit (Qiagen) before the 12 PCR cycle amplification step using the FailSafe PCR enzyme kit (Epicenter, WI) and selected ScriptSeq Index primers as reverse primers. The Agencourt AMPure XP system (BeckmanCoulter, NJ) purified the libraries and thereby size selected for > 200 bp. Each library’s size and quality was assessed on the Agilent 2100 Bioanalyzer and a High Sensitivity DNA Chip (Agilent Technologies, Wilmington, DE) and quantified with the NEBNext Library Quant Kit Protocol (New England BioLabs). The NextSeq 500/550 High Output Kit (300 cycles) was used for sequencing using the The Illumina NextSeq 500.

Sequences were initially analyzed using CLC Genomics software, version 6.0. Sequences with a quality score of less than 30 were discarded, the remaining sequences were trimmed, and sequences of less than 36 bp were discarded. Sequences were mapped to the *E. coli* W3110 genome, (NCBI accession number NC_007779.1) using the following CLC genomics mapping parameters: mismatch, 1; insertion, 3; deletion, 3; length, 0.9; similarity, 0.95; auto-detect paired distances on and map randomly. CLC RNA-seq was performed using the following parameters: mismatch, 2; length fraction, 0.9; similarity fraction, 0.95; strand specific selected; maximum 3 hits, 3; paired settings, 36 to 500; broken pairs counting selected. Only unique counts generated for individual genes were used as the starting data for all subsequent analyses.

Differential expression analysis was performed using the R package DESeq2. Reported log-fold-changes represent difference in expression of each gene in the evolved strains in 5 mM benzoate relative to the ancestor in the same condition. We also performed a control comparing the ancestor in 5 mM benzoate to the ancestor without benzoate. A gene was said to be differentially expressed if it had a log-fold-change greater than 1, and p-value < 0.01.

### Accession numbers for resequenced genomes and for RNAseq

The SRA accession number for RNAseq files is: PRJNA491479.

## ACKNOWLEDGMENTS

This work was supported by award MCB-1613278 from the National Science Foundation, and by Summer Science funds from Kenyon College.

